# Electronic blood vessel

**DOI:** 10.1101/2020.05.19.103457

**Authors:** Shiyu Cheng, Chen Hang, Li Ding, Liujun Jia, Lixue Tang, Lei Mou, Jie Qi, Ruihua Dong, Wenfu Zheng, Yan Zhang, Xingyu Jiang

## Abstract

Advances in bioelectronics have great potential to address unsolvable biomedical problems in the cardiovascular system. By using poly(L-lactide-co-ε-caprolactone) (PLC) that encapsulates the liquid metal to make flexible and bio-degradable electrical circuitry, we develop an electronic blood vessel that can integrate flexible electronics with three layers of blood vessel cells, to mimic and go beyond the natural blood vessel. It can improve the endothelization process through electrical stimulation and can enable controlled gene delivery into specific part of the blood vessel via electroporation. The electronic blood vessel has excellent biocompatibility in the vascular system and shows great patency three months post-implantation in a rabbit model. The electronic blood vessel would be an ideal platform to enable diagnostics and treatments in the cardiovascular system and can greatly empower personalized medicine by creating a direct link of vascular tissue-machine interface.

## Introduction

Cardiovascular diseases remain the number one cause of mortality worldwide^1^. In treating cardiovascular diseases via coronary artery bypass grafting (CABG) surgery, none existing small diameter (<6 mm) tissue engineered blood vessel (TEBV) has met clinical demands^2^. To fabricate the TEBV, a range of approaches, such as decellularized matrix^3–7^, self-assembly cell sheets^8–11^, bioactive and biomimetic materials^12–14^, were developed and clinically investigated in recent years. However, most of these methods only serve as scaffolds to provide mechanical support and mainly rely on the remodeling process by the host tissue and present significant limitations in helping the regeneration of neo blood vessel. Thus far, none of them have achieved satisfactory clinical results. Specifically, a complex interplay between the blood flow and the TEBV can often cause inflammatory responses, resulting in thrombosis, neointimal hyperplasia or smooth muscle cell accumulation nearby the scaffold^2,15^, in different pathological stages. To address these issues, next-generation TEBV should not only function as scaffolds to provide the mechanical support and facilitate host cell recruitments, but also have the capability to actively respond to and couple with the native remodeling process in order to provide adaptive treatments after implantation.

Combining living tissues with flexible electronics^16,17^ could endow the conventional TEBV more functionalities and capabilities to overcome existing biomedical problems, such as precision diagnostics, by *in situ* sensing the blood flow and temperature, and treatments by therapeutic drug or gene delivery^18,19^. In previous works, we have developed many approaches to fabricate structures that mimic natural blood vessel with multiple types of blood vessel cells in different layers, including stress-induced self-rolling membrane^13,20–22^ and layer by layer technique^23^. Recently, we have developed a printable metal-polymer conductor (MPC) which exhibited excellent compatibility and high stretchability^24,25^. Here, we develop an electronic blood vessel that integrates flexible electrodes into the bio-degradable scaffold by combining the liquid metal with poly(L-lactide-co-ε-caprolactone) (PLC) into a metal-polymer conductor. As a proof of concept, we used the electronic blood vessel to carry out *in vitro* electrical stimulation and electroporation. By electrical stimulation, the electronic blood vessel can effectively promote cell proliferation and migration in a wound healing model. It can also *in situ* deliver the GFP DNA plasmid into three kinds of blood vessel cells via electroporation. We evaluated the efficacy and biosafety of the electronic blood vessel in the vascular system through a three month *in vivo* study by using a rabbit carotid artery replacement model and confirmed its patency by using ultrasound imaging and arteriography. Our results pave the way to integrating flexible, degradable bio-electronics into the vascular system, which can serve as a platform to carry out further treatments, such as gene therapies, electrical stimulation, electronically controlled drug releases.

## Results

### Fabrication of the electronic blood vessel

We fabricated the electronic blood vessel (**Figure 1A**) by rolling up a PLC based metal-polymer conductor (MPC-PLC) membrane (**Figure 1B**) with the assistance of a polytetrafluoroethylene (PTFE) mandrel. The MPC circuit was well distributed in the three dimensional (3D) multilayered tubular structure. The inner diameter of the electronic blood vessel was around 2 mm **(Figure 2A)** and the minimum diameter could be around 0.5 mm (**Figure S1A**). The MPC-PLC membrane is flexible and degradable and the MPC circuit is conductive (**Figure 1B-D**). The conductivity of the MPC circuit is about 8*10^3^ S cm^−1^ and the ΔR/R of the circuit remained constant after around 1000 cycles of bending and rubbing (**Figure 1C**). The PLC is projected to be entirely degraded around 1~2 years by the manufacturer. The MPC-PLC membrane underwent a mass loss of around 10% during 8-week incubation in the phosphate-buffered saline (PBS, 37 °C) (**Figure 1D**). We observed a relatively quick degradation in the first week (**Figure S2**). To fabricate the MPC-PLC membrane, we screen-printed the conductive ink on the polyethylene terephthalate (PET) membrane (**Figure 1E**). The electrode design was optimized for electroporation and electrical stimulation. By preparing different electrode designs, we could either target individual blood vessel layers (tunica intima/media/adventitia) with the electrodes distributed in specific areas or target all three layers with the full electrode (**Figure S1B-C**). We prepared the liquid metal conductive ink by sonicating the mixture of Gallium-Indium alloy (EGAIn, ≥99.99%, Sigma, US) and the volatile solvent (1-decanol, Macklin, Shanghai) (**Figure 1F, H**). The liquid metal particles (LMPs) exhibit a core-shell structure, where the core is the Ga-In alloy and the shell is the Ga-In Oxide (**Figure 1G**). The diameter of the LMPs is around 2 μm (**Figure 1J**). We embedded the LMPs-based circuit with the PLC solution (5 wt % in CH_2_Cl_2_) and peeled the MPC-PLC membrane off the PET substrate after evaporation of the solvent in a chemical hood (**Figure 1E**). The thickness of the MPC-PLC membrane is about 50 μm and it is tunable by changing the volume of the PLC solution. The LMPs could be broken during the process of peeling and release the Ga-In alloy to make the circuit conductive^25^. We confirmed the structure of the conductive circuit via corrosion of the Ga-In alloy by adding excessive hydrochloric acid (HCl, Beijing Chemical Works, China). The LMPs were evenly distributed in the cellular PLC host (**Figure 1K, L**). The liquid metal could form a coherent conductive pathway in the polymeric host.

**Figure 1.**
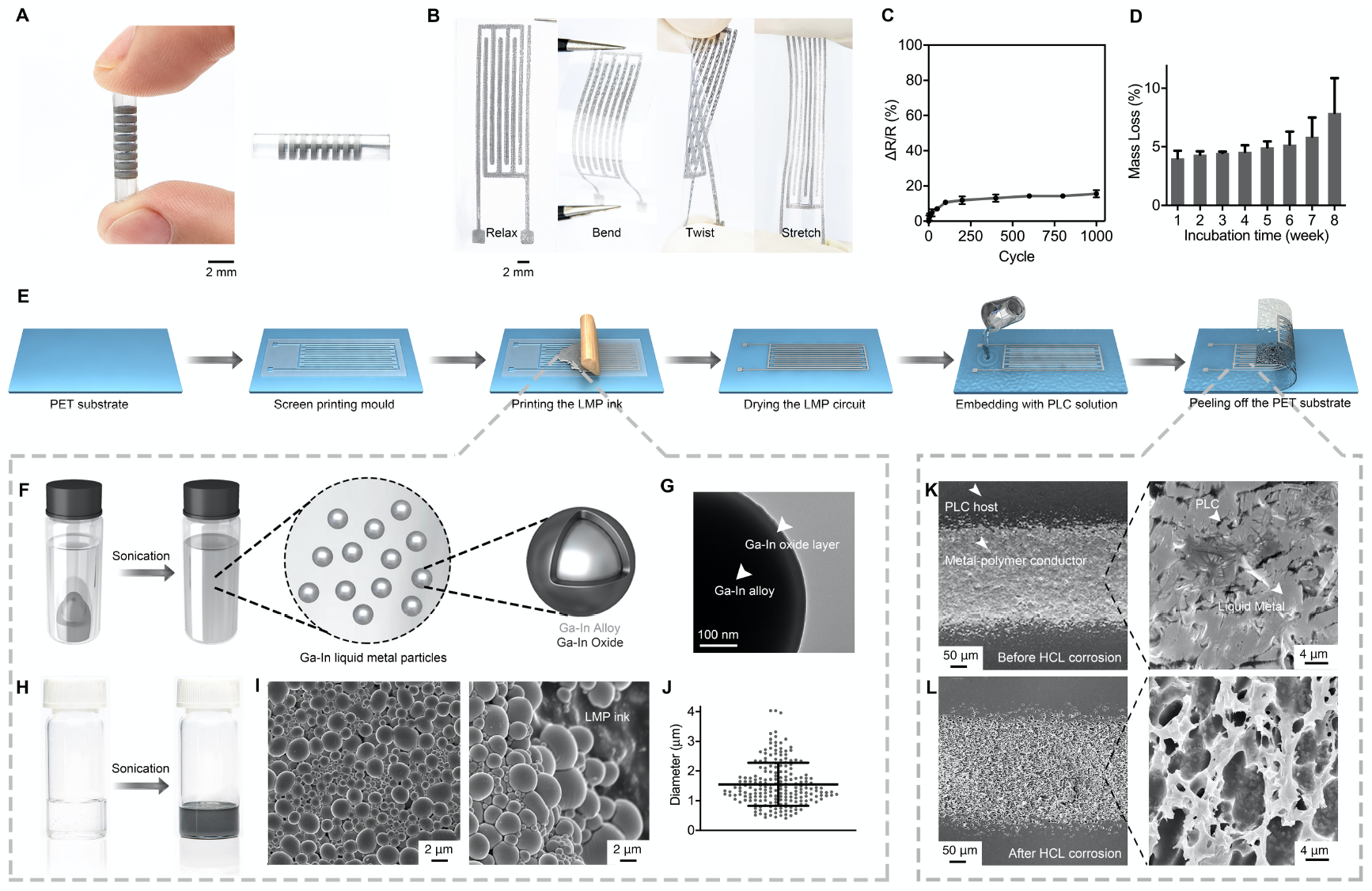
Fabrication and characterization of the electronic blood vessel. (**A)** Snapshots of the electronic blood vessel. Scale bar, 4 mm. **(B)** Snapshots of the MPC-PLC membrane. Scale bar, 1 mm. **(C)** ΔR/R changes with a bend of 180° for 1000 cycles (n=3). **(D)** Mass loss of the MPC-PLC membrane during 8-week incubation (n=3 for each time point). **(E)** Schematic of fabrication of the MPC-PLC membrane. **(F)** Schematic of fabrication of LMPs. **(G)** TEM image of the Ga-In particle. **(H)** Photographs of mixture of the Ga-In alloy and the solvent, and the LMP ink after sonication. **(I)** Representative SEM images of LMPs. Scale bar, 2 μm. **(J)** Diameter distribution of LMPs (n =200). **(K)** SEM images of the MPC-PLC circuit. White arrowheads indicate the PLC host and MPC-PLC membrane. Scale bar, 50 μm. Right is the Zoom-in view. White arrowheads indicate the PLC and LMP ink. LMPs were embedded by the PLC as the host polymer. Scale bar, 4 μm. **(L)** SEM images of the MPC-PLC circuit after corrosion by excessive hydrochloric acid. Scale bar, 50 μm. Right is the Zoom-in view. LMPs were fully removed by corrosion. The porous structure only consisted of the PLC. Scale bar, 4 μm.

**Figure 2.**
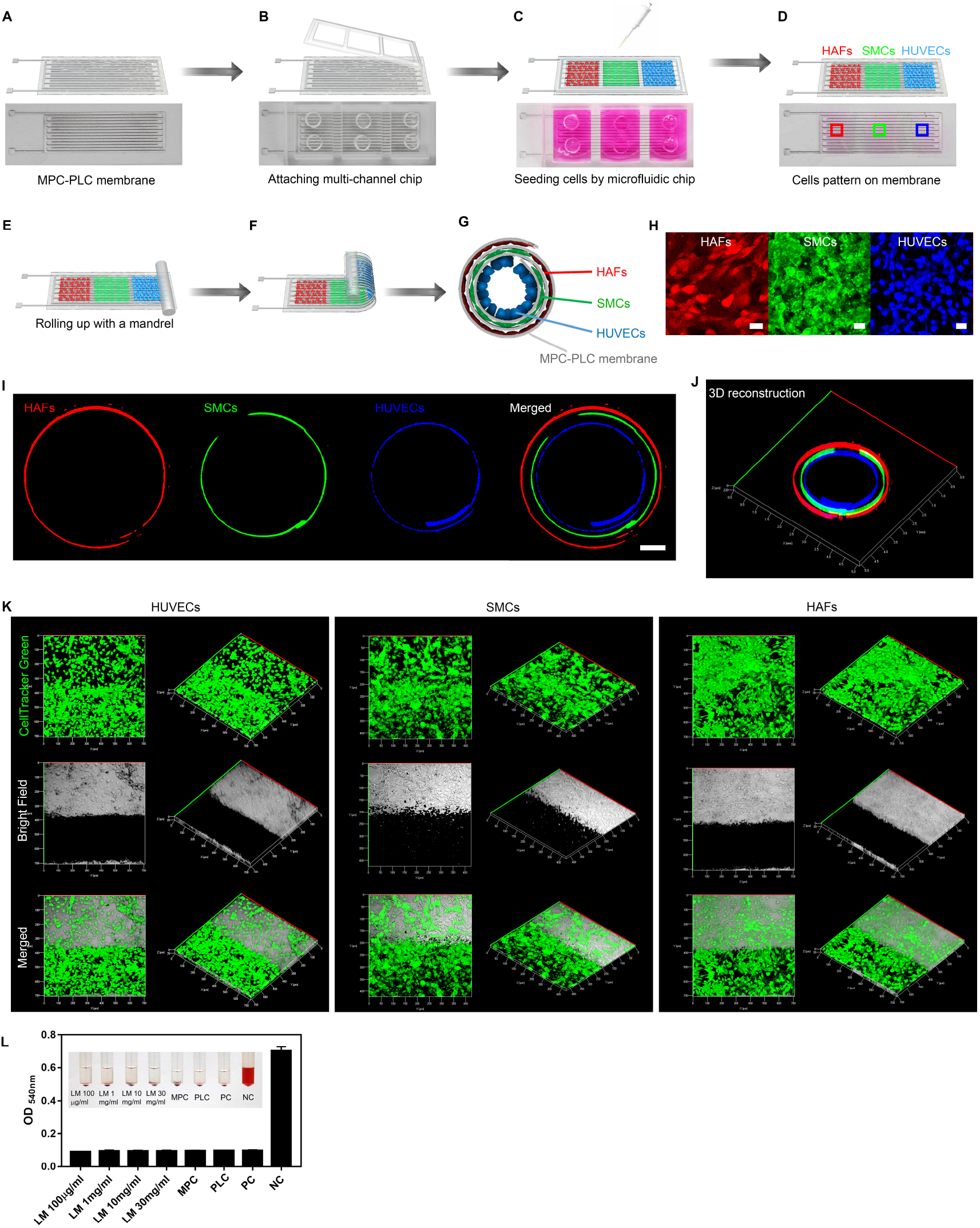
*In vitro* biocompatibility of the electronic blood vessel. **(A)** The MPC-PLC membrane. **(B)** Attaching a multichannel microfluidic chip onto the MPC-PLC membrane. **(C, D)** Patterning three kinds of blood vessel cells (blue: HUVECs, green: SMCs, red: HAFs) onto the MPC-PLC membrane by using PDMS microfluidic chip. **(E, F)** Rolling the cell-laden membrane into a multi-layered tubular structure with a PTFE mandrel. **(G)** Cross-sectional view of natural blood vessel mimicking structure with different cells distributed in different layers. **(H)** Fluorescent images of cells on the MPC-PLC membrane, in corresponding to (D). Stained by CellTracker violet, green, deep red, respectively. Scale bar, 20 μm. **(I)** Three kinds of blood vessel cells distributed in different layers of the electronic blood vessel. Scale bar, 500 μm. **(J)** 3D construction of (I). **(K)** Cell viability after 2-week incubation in the electronic blood vessel. Stained by CellTracker green. The right column is the 3D reconstruction of the left column in each image. The black area represents the MPC circuit and the transparent area represents the PLC in bright-field images. **(L)** Hemolysis test. The MPC circuit, PLC membrane, and liquid metal with different concentrations were tested with the rabbit whole blood and exhibited good blood compatibility. Saline was used as the positive control whereas water was used as the negative control.

### *In vitro* characterization of the electronic blood vessel

To evaluate biocompatibility of the electronic blood vessel, we used the microfluidic technology to realize accurate 3D pattern of three kinds of blood vessel cells in a natural blood vessel mimicking fashion. By employing a multichannel microfluidic chip, we delivered human umbilical vein endothelial cells (HUVECs, blue), human aortic smooth muscle cells (SMCs, green), human aortic fibroblasts (HAFs, red) sequentially on the MPC-PLC membrane (**Figure 2A-E**). We designed the width of each channel to match the circumference of each layer of the tube based on thickness of the MPC-PLC membrane and the diameter of the tube (**Figure 2B**). To distinguish different cell types, we stained the HUVECs, SMCs, HAFs with different fluorescent dyes (HUVECs: Cell Tracker Violet; SMCs: Cell Tracker Green; HAFs: Cell Tracker Deep red) before seeding them into the microfluidic chip. After 1 d incubation in culture medium (Dulbecco’s modified eagle medium, DMEM, 10% FBS, 37 °C, 5% CO_2_), cells attached on the MPC-PLC membrane (**Figure 2D, H**). We peeled the microfluidic chip off the cell-laden MPC-PLC membrane and rolled it up with a PTFE mandrel, forming a 3D multi-layered tubular structure (**Figure 2E-G**) with HUVECs, SMCs, and HAFs evenly distributed in the inner layer, middle layer, and outer layer, respectively (**Figure 2I, J**). This structure well mimics the structure of the natural blood vessel. To better understand the blood vessel cells distributed in different layers of the electronic blood vessel, we stained HUVECs, SMCs, HAFs, and the MPC-PLC layer with CellTracker DiO, CellTracker DiI, CellTracker DiD, and CellTracker Blue to show the relative distribution of different layers (**Figure S3**). We used a biomedical fibrin glue to facilitate the combination of different layers^18^. We incubated the cell-laden electronic blood vessel for 14 d and stained the cells with the Calcein AM green. The evenly distributed green color on the MPC-PLC membrane indicated that the cells exhibited high viability after a 14 d culture (**Figure 2K**). We measured the transport of ions through the MPC-PLC membrane to further evaluate and quantify the permeability of the electronic blood vessel **(Figure S4)**. Ca^2+^, Fe^3+^, Mg^2+^ could permeate the electronic blood vessel over time. We also conducted a hemolysis test, which showed that the electronic blood vessel exhibited very good blood biocompatibility (**Figure 2L**). The *in vitro* characterization demonstrated that the electronic blood vessel exhibited excellent biocompatibility and we then conducted further functional tests of the embedded MPC circuits, i.e. electrical stimulation and electroporation.

### *In vitro* electrical stimulation promotes HUVEC proliferation and migration

To prove the functionality of the electronic blood vessel, we carried out *in vitro* electrical stimulation to improve proliferation and migration of HUVECs. Direct current (DC) electric field has been shown to effectively increase the angiogenesis *in vitro* and *in vivo*^26^. We patterned HUVECs on the MPC-PLC membrane (DMEM, 10% FBS, 37 °C, 5% CO_2_) by using a multi-chamber PDMS chip. The initial number of cells in each chamber was the same (3×10^4^). After 12 h incubation, we applied different DC voltages to yield different electrical field strength: 25, 50, 75 100, 200, and 400 mV mm^−1^, respectively (**Figure 3A, B**). After 2 d incubation and electrical stimulation, we randomly selected 6 different domains on each sample and analyzed by using a laser scanning confocal microscopy (LSCM, LSM 710, Zeiss, Germany). We stained nuclei (blue) with Hoechst 33342 (Invitrogen, USA) and stained the living and dead cells with Calcein-AM (green) and PI (red), respectively. The green cells occupied the whole domain, indicating that electrical stimulation did not hurt proliferation of HUVECs (**Figure 3C**). We counted the number of nuclei by using ImageJ. The cell number under 50 mV mm^−1^ was highest, about 2.4 times that of the control (**Figure 3E**). We used the CCK-8 kit to confirm this conclusion (**Figure 3F**). We speculated that the DC electric field had selectively regulated production of certain growth factors and cytokines important for angiogenesis.

**Figure 3.**
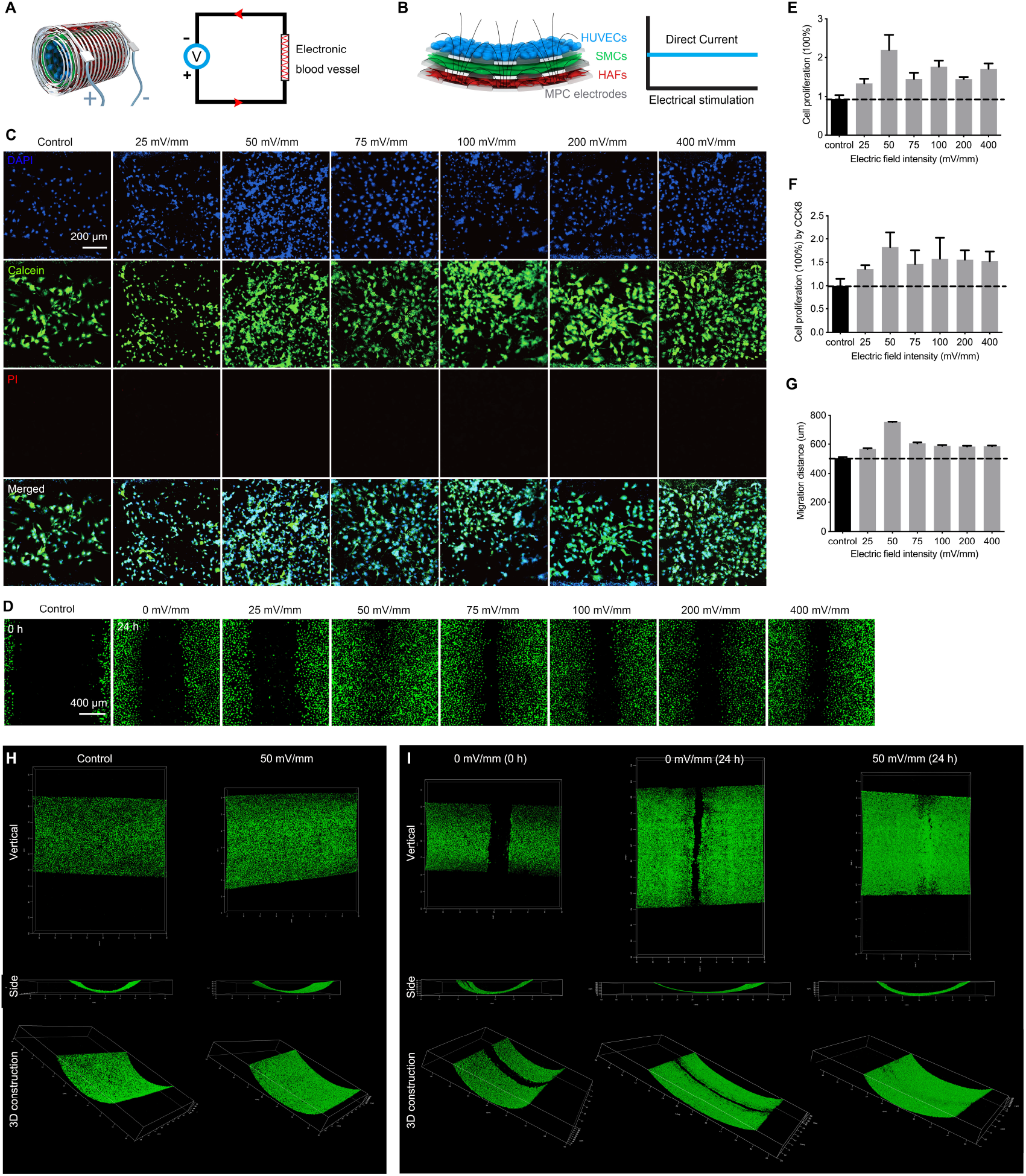
*In vitro* electrical stimulation. **(A-B)** The electronic blood vessel was connected to electrochemical station (A) to generate a direct current voltage (B). **(C)** Confocal fluorescent images of HUVEC proliferation after 2 d incubation and stimulation under different DC electric field: 25, 50, 75, 100, 200, 400 mV/mm, respectively. Blue, DAPI; green, Calcein AM; red, PI. **(D)** Confocal fluorescent images of HUVEC migration after 24 h incubation and stimulation under different DC electric field: 25, 50, 75, 100, 200, 400 mV/mm, respectively. Stained by Calcein AM. Scale bar, 400 μm. A 10 μl tip was used to scratch a line to create a model for HUVECs migration. **(E)** The proliferation of HUVECs under different DC electrical field, setting the control (without electric field) as 100%. (n=4). **(F)** The proliferation of HUVECs under different DC electrical field tested with the CCK-8 kit. (n=4). **(G)** The migration of HUVECs at different DC electrical field (n=4). **(H)** The proliferation of HUVECs in a 3D model under DC electric field of 50 mV/mm. Stained by Calcein AM. **(I)** The migration of HUVECs in a 3D model under DC electric field of 50 mV/mm. Stained by Calcein AM. The HUVECs formed a complete endothelial layer after 24 h electrical stimulation.

We explored the migration of HUVECs under different DC electric field strengths. We made a scratch on a PDMS substrate by using a 10 μl tip. After application of an electric field of 50 mV mm^−1^, HUVECs migrated 750 μm and the wound completely healed after 24 h (**Figure 3D**). Different strengths had enhanced migration differently compared to the control group without electrical stimulation (**Figure 3G**). The *in vitro* DC electrical stimulation thus proved to have effectively promoted the proliferation and migration of HUVECs.

We further evaluated the effectiveness of electrical stimulation in a 3D model of endothelization. We patterned the HUVECs on the MPC-PLC membrane and made a scratch on the cell-laden membrane by using a 10 μl tip. We transformed the 2D cell-laden membrane into 3D cell-laden structure and connected it to the electrochemical station to test the proliferation and migration. We applied an electric field of 50 mV mm^−1^ on multiple samples and we observed the proliferation and migration at different time points. We stained the living and dead cells with Calcein-AM (green) and PI (red). We counted the cell number and the cell density was higher than the control group (Figure 3H). The HUVECs formed a complete endothelial layer after 24 h (Figure 3I). To better evaluate the biocompatibility of the MPC circuit under electrical stimulation, we extended the electrical stimulation time to 10 days and the live/dead staining showed that cells exhibited excellent viability (Figure S5).

### *In vitro* electroporation

To further prove the functionality of the electronic blood vessel, we designed multiple circuit patterns for electroporation, being able to target different pathological issues in different layers of blood vessel cells (**Figure S1B**). We conducted the electroporation with a circuit pattern that could target all three layers. We seeded the cells onto the MPC-PLC membrane and transformed it into 3D tubular structure for electroporation. We immersed the 3D cell-laden electronic blood vessel in the GFP plasmid DNA solution for 10 min before electroporation. The GFP plasmid DNA could also be lyophilized onto the MPC-PLC membrane before seeding cells (**Figure S7A**) and transforming to 3D structure for electroporation. We connected the 3D cell-laden electronic blood vessel with an electroporator (BTX, CM630, US) to generate DC pulses (**Figure 4A**) and achieved the delivery of the green fluorescent protein (GFP) DNA plasmid in three kinds of blood vessel cells (**Figure 4B**). To optimize the performance of the electronic blood vessel, we found two major parameters determining the efficacy of electroporation, including voltage and pulse duration. We conducted the electroporation with different voltages (40 V/60 V/80 V) and pulse durations (100 μs/1 ms). If the voltage was too low, it would cause low efficacy or no transfection; if the voltage was too high or if the pulse duration was too long, it would cause low efficacy and cell death (**Figure S6**). To realize the optimal efficacy, we exerted a square wave with the voltage of 60 V, pulse duration of 100 μs, and pulse interval of 1 s for 5 times. We delivered GFP plasmid DNA into three kinds of blood vessel cells and the GFP DNA realized expression with more than 95% of cells showing green fluorescence (**Figure 4B**). We observed successful expression of GFP among all three layers of the blood vessel cells and exhibited a uniform 3D distribution in the electronic blood vessel **(Figure 4C)**. To evaluate the potential of the electronic blood vessel for *in vivo* electroporation, we lyophilized the GFP plasmids DNA on the MPC-PLC membrane to test its effectiveness **(Figure S7A)**. We carried out electroporation by attaching the plasmids-laden MPC-PLC membrane to an isolated rabbit vascular tissue with the voltage of 60 V, pulse duration of 100 μs, and pulse interval of 1 s for 5 times. We observed successful expression of GFP in the isolated rabbit vascular tissue **(Figure S7A)** after 2 d incubation. These promising *in vitro* results encouraged us to carry out the *in vivo* tests of the electronic blood vessel.

**Figure 4.**
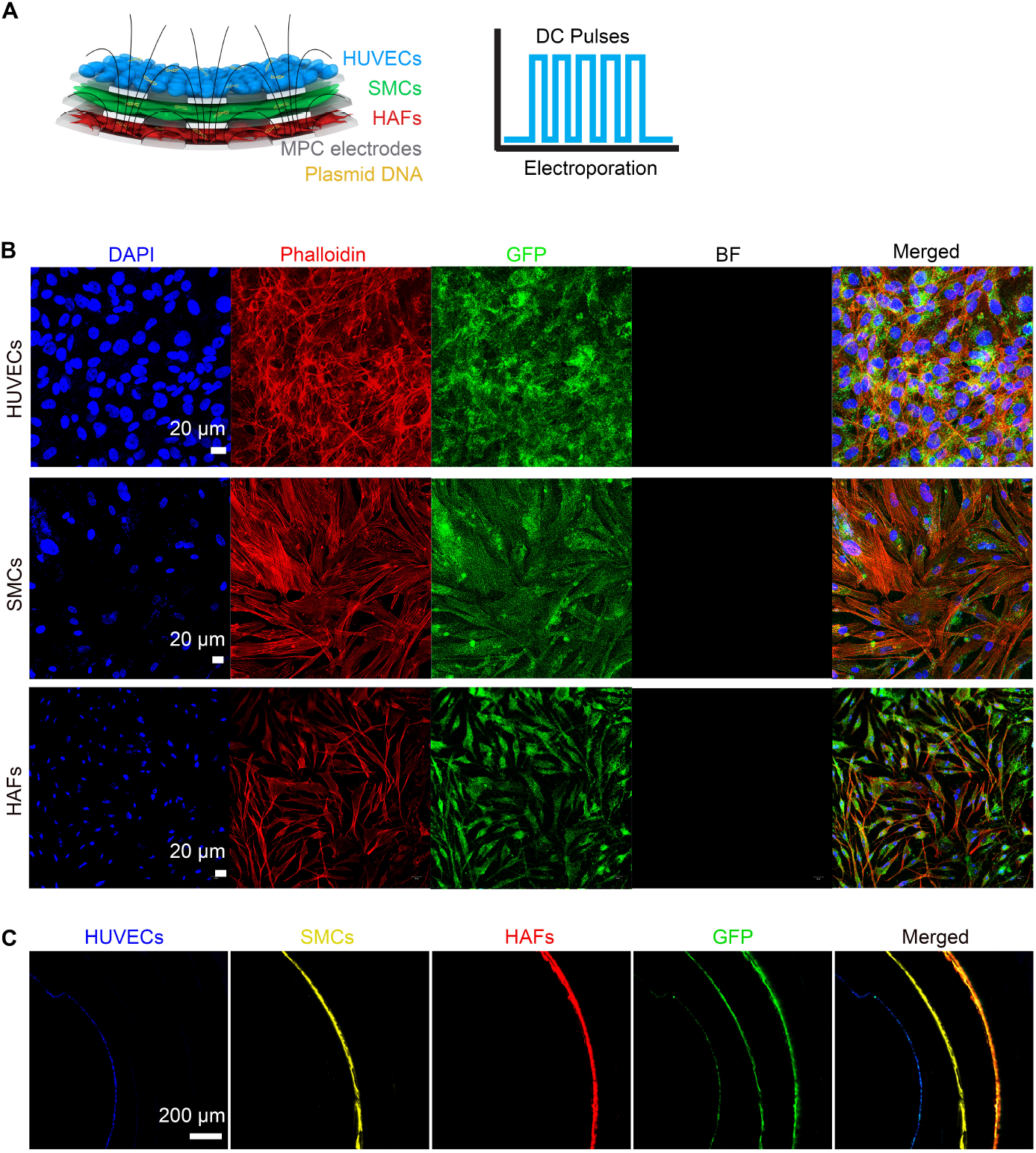
*In vitro* electroporation. **(A)** The electronic blood vessel was connected to an electroporator to generate electrical pulses. **(B)** Confocal fluorescent images of HUVECs, SMCs, and HAFs after electroporation under condition: voltage 60 V; pulse duration 100 μs; 5 pulses; pulse interval 1 s. Upper: HUVECs; middle: HAFs; lower: SMCs. Blue represents cell nucleus, stained by DAPI; red represents cell skeleton, stained by Rhodamine Phalloidin; green represents GFP; black represents that cells were on the top of non-transparent MPC-PLC circuit, bright field. Scale bar, 20 μm. **(C)** 3D distribution of HUVECs, SMCs, HAFs after electroporation under condition: voltage 60 V; pulse duration 100 μs; 5 pulses; pulse interval 1 s. Blue represents HUVECs, stained by CellTracker Blue; yellow represents SMCs, stained by CellTracker DiI; red represents HAFs, stained by CellTracker DiD; green represents green fluorescent protein. Scale bar, 200 μm.

### Mechanical properties of the electronic blood vessel

To find out whether the electronic blood vessel is suitable for *in vivo* implantation, we measured the mechanical properties, including stress-strain curve, compliance, and burst pressure, of the electronic blood vessel with a diameter of 2 mm prior to implantation (**Figure 5A-F and Figure S8**). The elastic modulus of the electronic blood vessel is about 130 MPa, the value is much higher than the native carotid artery. The ultimate tensile strength of the electronic blood vessel is about 27 MPa, the value is also much higher than that of the native carotid artery. The initial compliance of the electronic blood vessel (n= 5) is about 5% per 100 mmHg in the range of 80-120 mmHg, the value is apparently below that of the native carotid artery (n= 3). The burst pressure of the electronic blood vessel (n= 5) is about 2800 mmHg, the value is similar to that of the native carotid artery (n= 3). The elongation at break of the electronic blood vessel is about 650% (n= 5), which is the twice as the value of native carotid artery (n= 3). The mechanical property of electronic blood vessel was considered robust enough for implantation.

**Figure 5.**
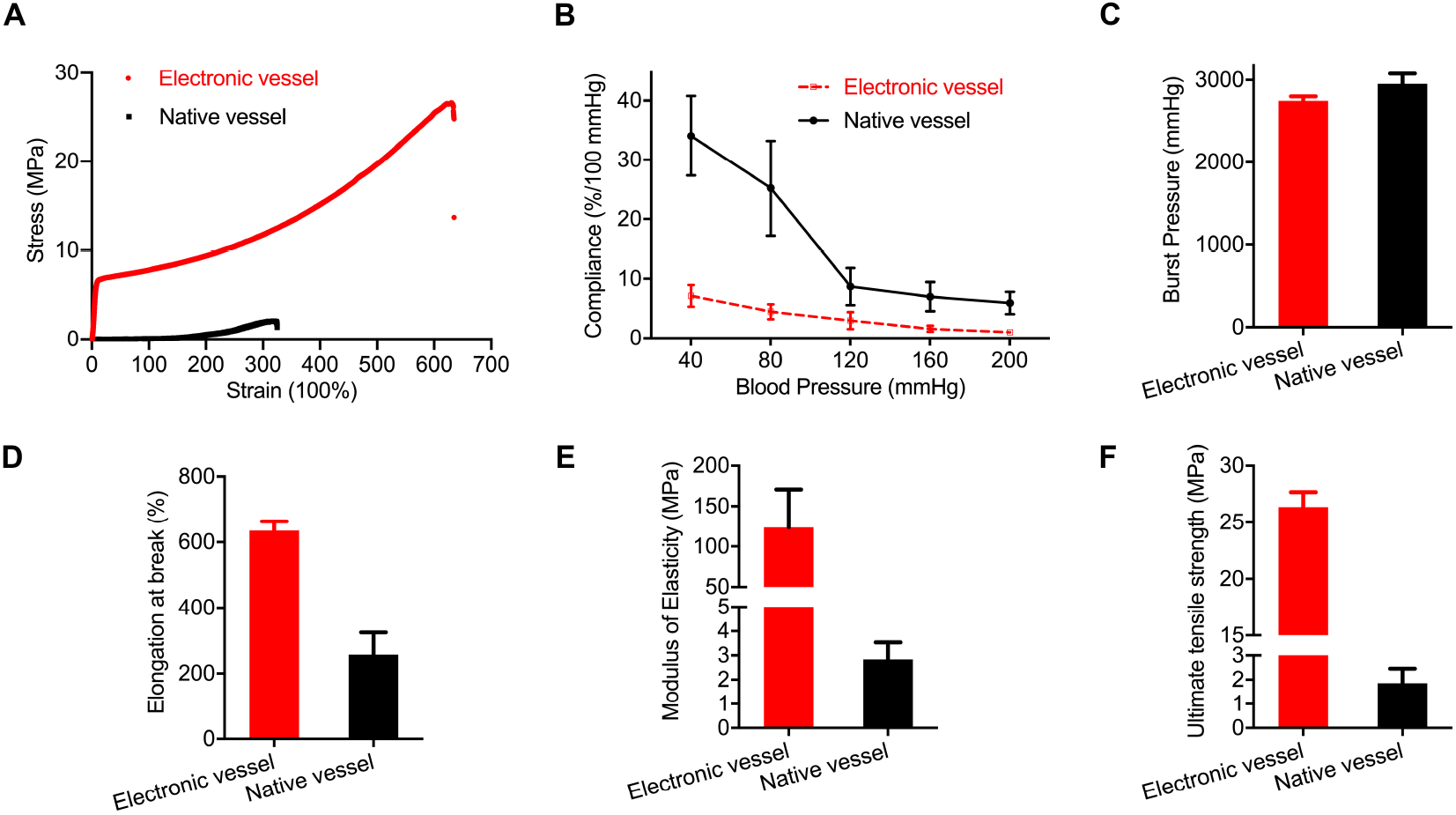
Mechanical properties of the electronic blood vessel. **(A)** Stress-strain curve. **(B)** Compliance test. **(C)** Burst pressure test. **(D)** Elongation at break. **(E)** Modulus of elasticity. **(F)** Ultimate tensile strength. Electronic blood vessel (n= 5) and native carotid artery (n= 3) were used in each test. All data are expressed as mean ± SD.

### *In vivo* implantation in rabbits and *in situ* monitoring

To investigate the electronic blood vessel as a vascular implant, we chose the New Zealand rabbit (age: 200-300 days, body weight: 3-4 kg) as the animal model and replaced the native carotid artery with the electronic blood vessel (**Figure 6A-C**). To avoid possible immunological response of the host tissue, we used the acellular electronic blood vessel in the preliminary *in vivo* study. We *in situ* monitored the implanted electronic blood vessel by doppler ultrasound imaging (**Figure 6D-I**) and the arteriography (**Figure 6J, K**). Doppler ultrasound imaging showed that the electronic blood vessel allowed for good blood flow 3 months post-implantation (**Figure 6D-G and Video S1**). The asymmetric velocity curve synchronized with the ultrasonic pulses indicated that the signal is from the carotid artery rather than the vein (**Figure 6D**). The diameter of the electronic blood vessel remained at a relatively constant value, about 2.3 mm, during the half month to three months post-implantation (**Figure 6G, H**). The mean velocity of the blood flow in different samples at different time point was about 0.47 m s^−1^, which was in the range of the normal value (**Figure 6I**). As the gold standard of the blood vessel patency, arteriography showed that the electronic blood vessel matched with the native carotid artery very well and allowed for excellent blood flow (**Figure 6J, K, Figure S9, and Video S2**). There was no sign of narrowing. The electronic blood vessel allows straightforward visualization under arteriography because the liquid metal-based circuitry has sufficiently high contrast over host tissues (**Figure 6J**). The red frame outlined the electronic blood vessel with an alternate strip structure from the MPC circuit.

**Figure 6.**
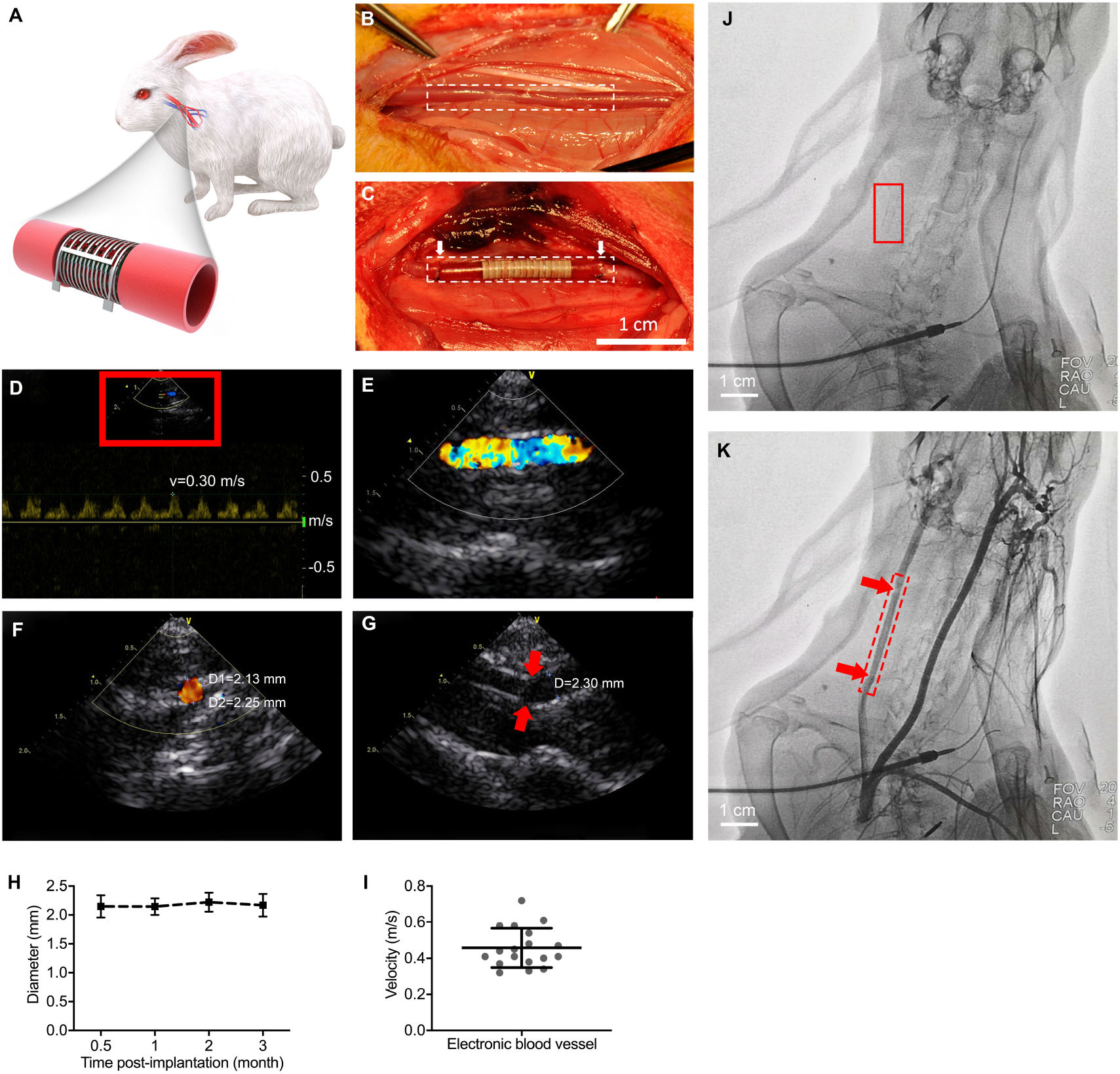
*In situ* monitoring of the electronic blood vessel. **(A)** Schematic of the electronic blood vessel in the carotid artery of the rabbit. **(B, C)** An end-to-end anastomosis procedure of electronic blood vessel implantation on carotid arteries of rabbits (n=6). The dotted frames outline the margin of the native carotid artery and the implanted electronic blood vessel. The white arrows indicate two ends of the electronic blood vessel. Scale bar, 1 cm. **(D-G)** *In situ* monitoring of the electronic blood vessel by doppler ultrasound imaging 3 months post-implantation. Representative images from at least three different animals. **(D, E)**The real-time blood flow at the operational site and the synchronized ultrasound pulses. The asymmetric velocity curve represents the signal is from carotid artery rather than vein. **(E)** Zoom-in view of red box in (D). **(F)** The cross view of the blood flow. **(G)** The suture site (red arrows) connecting the native carotid artery and the electronic blood vessel. **(H)** The diameter changes of the electronic blood vessel in different time post-implantation. **(I)**The velocity of blood flow at the operational site (n=18, from different rabbits at different time points). **(J, K)** *In situ* monitoring by arteriography 3 months post-implantation. **(J)** Image before injecting the contrast media. Red box indicates the position of the implanted electronic blood vessel. **(K)** Image after injecting the contrast media. Red box indicates the position of the implanted blood vessel. Red arrows indicate the suture sites connecting the native carotid artery and the electronic blood vessel.

### *Ex vivo* study

We dissected all the implanted electronic blood vessels three months post-implantation for characterization. The lumen and the outer surfaces of the explanted electronic blood vessel were smooth, covered by the remodeling tissues (**Figure 7A, B**). The diameter of the native blood vessel significantly reduced due to the lack of the blood pressure, whereas the electronic blood vessel remained the same as before (**Figure 7C**). We observed the micro structure of the circuit by using the SEM after explanting the electronic blood vessel from rabbit. The MPC-PLC membrane still maintained interdigitated structure with the MPC circuit and PLC host (**Figure 7E**). There was a layer of neo-tissue formed, which well covered the MPC-PLC membrane (**Figure 7F-H**). We could also see some red blood cells on the top of the circuits, which were similar in number to those on native blood vessels (**Figure 7D**). We tested the conductivity of the circuit in the electronic blood vessel. The MPC circuit was still conductive and the conductivity was around 7.2 *10^3^ S cm^−1^.

**Figure 7.**
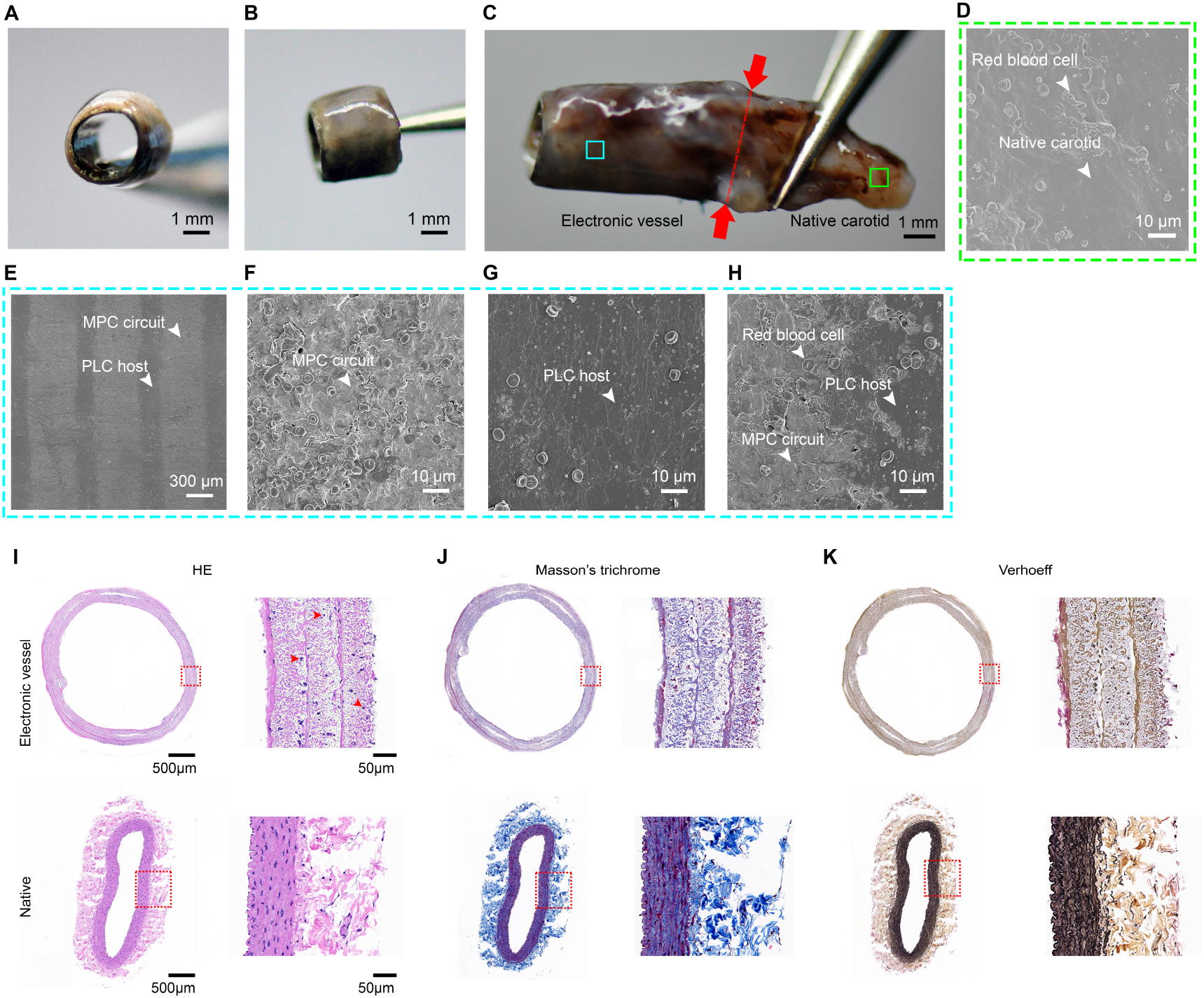
*Ex vivo* study of the electronic blood vessel after 3 months implantation in the rabbit. **(A-C)** The cross-sectional (A) and lateral view (B, C) of the explanted electronic blood vessel after 3-month host remodeling. Red arrows and the dotted line indicate the suture site connecting the native carotid artery (right) and the electronic blood vessel (left). Scale bar, 1 mm. **(D)** Representative SEM image of the lumen of the native carotid. Scale bar, 10 μm. (**E**) Representative SEM image of the lumen of the electronic blood vessel. Scale bar, 300 μm. (**F-H**) Zoom-in view of the MPC circuit (F), PLC host (G) and the interconnection area of MPC circuit and PLC host (H). Scale bar, 10 μm. **(I-K)** Hematoxylin/eosin (H&E) staining, Masson’s trichrome staining, and Verhoeff’s staining of the implanted electronic blood vessel 3 months post-implantation, setting native carotid artery as control. The right panel is the zoom-in view of the left panel. Scale bar (left), 500 μm. Scale bar (right), 50 μm.

To study the histological changes of the implanted electronic blood vessels, we performed histological staining of the electronic blood vessels, setting native carotid blood vessel as a positive control. Hematoxylin/eosin (H&E) staining (**Figure 7I**) of the cross section of the electronic blood vessel is round-shaped, continuous, red, which is similar as the native blood vessel. The three-layered MPC-PLC membrane merged into one intact layer with secretion of substantial extracellular matrix between the different layers of electronic blood vessel. We could clearly see dark blue nuclei (red arrows in Figure 7I) in all the layers, which indicated successful migration and infiltration of host cells into the electronic blood vessel. We compared the cell density in the electronic blood vessel with the native carotid. The cell density of the electronic blood vessel was around 400 cells mm^−2^ whereas that of the native carotid was around 535 cells mm^−2^. More importantly, a dense layer of cells with curved structure were formed in the lumen of the electronic blood vessel, which indicated the excellent endothelization and thus ensured good blood flow. To further confirm the components in the implanted electronic blood vessel, we performed Masson’s trichrome (**Figure 7J**) and Verhoeff’s staining (**Figure 7K**). Masson’s trichrome staining can stain and assess keratin and muscle fibers (red) or collagen (blue) and Verhoeff’s staining can stain and assess the presence of elastin fibers. Masson’s trichrome and Verhoeff’s staining showed well-distributed collagen and elastin fibers both inside the layers and between different layers, indicating appropriate remodeling. Compared to the abundance level of extracellular matrix in the native blood vessel, the electronic blood vessel still requires more time for material degradation and tissue remodeling. These results indicated that the electronic blood vessel might still be functioning with the presence of conductive materials and host remodeling 3 months post-implantation.

To investigate the influence of the electronic blood vessel on the host, we performed the cross section and histological staining of the major organs, including heart, liver, spleen, lung, and kidney, together with the dissection of the implanted electronic blood vessel 3 months post-implantation. The H&E staining and Masson’s Chrome staining showed that there was no significant pathological changes or inflammatory responses in these organs (**Figure S10**). To evaluate whether there were chronic inflammation or infection, we conducted the ELISA assay of three important proteins in the blood, including interleukin-6 (IL-6), procalcitonin (PCT), and C-reactive protein (CRP) (**Figure S11**). The concentrations of IL-6 and PCT were not higher than the normal value of healthy rabbits (red dotted lines). The concentration of CRP was higher than the normal value of healthy rabbits (red dotted line). All three indexes decreased over time, a tendency that was expected. The results showed that most of the indexes were in the normal range and there were no significant pathological changes or inflammatory responses. We also conducted the complete blood count of the rabbit, including white blood cell count, absolute neutrophil count, and absolute lymphocyte count, most of which were in the normal range over time (**Figure S12**). These results confirmed that as an implant for vascular system, the electronic blood vessel had no significant detriment to the host.

## Discussion

None of the existing small-diameter TEBVs has met the demands in treating cardiovascular diseases. Conventional TEBVs can be greatly improved to provide next-generation treatment by integrating with flexible bioelectronics. In this work, we report an electronic blood vessel with excellent biocompatibility, flexibility, mechanical strength, and degradability by combining the MPC with a US FDA-approved biodegradable polymer. As a proof of concept, we verified that the electronic blood vessel can accelerate the HUVEC proliferation and migration by electrical stimulation, thus facilitating the endothelization process, which is important to an engineered vascular conduit in preventing early thrombosis. Because most TEBVs occluded with 2 weeks post-implantation, our electronic blood vessel, with a patency of at least 12 weeks, is of great promise for clinical application. We also showcased that it could be used to perform *in situ* gene delivery via electroporation, which laid a foundation for future design and optimization such that we can carry out further gene therapies targeting different pathological problems after implantation.

The electronic blood vessel has a high level of safety. The liquid metal^16,27^ has been proven to be highly biocompatible and the PLC has been approved by the US FDA for implants^28^. We validated its biosafety in the vascular system by a 3-month implantation in a rabbit carotid artery model. Both the *in situ* monitoring, including doppler ultrasound imaging and arteriography, and *ex vivo* study demonstrated that it was safe as an electronic implant in the vascular system and at the body level. The electronic blood vessel exhibited higher strength than the native blood vessel. The potential harm of a rigid synthetic blood vessel is the mismatch with host tissue after implantation. However, from the in vivo results, these discrepancies did not bring about any major issues and the electronic blood vessel matched very well with the host carotid artery during the time of *in situ* monitoring (Figure 6J, K, Video S2). The reasons why we chose liquid metal in the electronic blood vessel are as following: i) comparing with gold or platinum, the Ga-In liquid metal allows superior flexibility and stretchability while maintains good conductivity, which is critical for an artificial blood vessel to adapt to the deformation due to the rhythmic beating; ii) it exhibits excellent cytocompatibility and blood compatibility according to our results; iii) compared to other sophisticated microfabrication techniques, using the screen printing technique is much more straightforward and could enable industrial-scale mass production in a cost-effective manner.

As a vascular substitute, the electronic blood vessel breaks through the limitations of the existing vascular scaffold by endowing the electrical function to conventional biodegradable TEBV and provides us a new platform for tackling the problems threatening the small-diameter blood vessel. By integrating with other electronic devices, the electronic blood vessel can provide various treatments, such as electrical stimulation, electroporation, electrically controlled drug release, and so forth. When combining with emerging technologies such as artificial intelligence, it can greatly boost future personalized medicine by bridging the vascular tissue-machine interface and empowering the health data collection and storage, such as blood velocity, blood pressure, and blood glucose level. In the future, optimizing its function and creating a multifunctional electronic blood vessel can greatly benefit human cardiovascular health.

## Conclusions

In this work, by integrating the liquid metal-based conducting circuitry with a biodegradable polymer, we develop an electronic blood vessel, with excellent biocompatibility, flexibility, conductivity, mechanical strength, and degradability, that enables *in situ* electrical stimulation to facilitate the endothelization process and electroporation to deliver genes in specific layers of blood vessel cells. It exhibited excellent patency and biosafety 3 months post-implantation in the vascular system of a rabbit model. In the future, the electronic blood vessel can be integrated with other electronic components and devices to enable diagnostic and therapeutic functions and greatly empower personalized medicine by creating a direct link of vascular tissue-machine interface.

## Supplemental Information

Supplemental Information can be found online.

## Acknowledgements

This study was supported by the National Key R&D Program of China (2018YFA0902600, 2017YFA0205901), the National Natural Science Foundation of China (21535001, 81730051, 21761142006, 51973045), the Chinese Academy of Sciences (QYZDJ-SSW-SLH039, 121D11KYSB20170026, XDA16020902), Shenzhen Bay Laboratory (SZBL2019062801004), the Tencent Foundation through the XPLORER PRIZE, Beijing Science and Technology Plan Project (Z191100007619053), Postgraduate Innovation Foundation of Peking Union Medical College (2019-1002-28) and Teaching Reform Foundation of Postgraduate Education in Peking Union Medical College (10023201900202). We thank Dr. Zewen Wei and Dr. Deyao Zhao for discussions on the electroporation experiment. We thank Mrs. Barbara Althaus at EPFL for her advice in writing the manuscript.

## Author Contributions

X. J. and S. C. conceived the idea. S. C. designed the experiments, fabricated the blood vessel, conducted the characterization, *in vitro* tests, mechanical tests, *in vivo* tests and *ex vivo* tests, analyzed the data, and wrote the manuscript. C. H. fabricated the blood vessel, performed *in vitro* tests, mechanical tests, and *in vivo* tests. L. D. performed the *in vivo* and *ex vivo* tests. S. C., C. H., and L. D. contributed equally. L. T. assisted with the preparation of the liquid metal particles, MPC-PLC membrane and the electroporation experiments. L. J. and Y. Z. performed the implantation of the electronic blood vessel, doppler ultrasound imaging and arteriography. L. M. conducted the ELISA assay. J. Q. and R. D. assisted the TEM characterization. W. Z. gave inputs on the *in vitro* tests and *ex vivo* tests. Y. Z. supervised the *in vivo* and *ex vivo* study. X. J. supervised the project, and revised the manuscript. All the authors took part in the discussion and writing.

## Data Availability

All data are available from the corresponding authors upon reasonable request.

