## Supplementary material for "Electronic blood vessel": Electronic blood vessel_SI_20200808.pdf

### Experimental Procedures

#### Preparation of the liquid metal conductive ink

We added 2.5 g EGaIn (Gallium Indium eutectic, 99.99%, Sigma-Aldrich) into 1 mL 1-Decanol (98%, MACKLIN, China). We sonicated the mixture for 1 min with the power of 300W by using a sonicator (Scientz, Scientz-IID, China). We obtained the solution of liquid metal particles of 2.5 g $ml^{-1}$ and we used it as the conductive ink for screen printing.

#### Preparation of MPC-PLC membrane

We used a screen printing equipment (Taobao, China) and employed PET membrane (Taobao, China) as the substrate for screen printing. We screen-printed the conductive liquid metal ink onto a screen printing template (Taobao, China) with desired pattern. We dissolved the poly (L-lactide-co-ε-caprolactone) (PLC) (70:30, RESOMER®, Evonik, Germany) particles in dichloromethane ($CH_2Cl_2$) at 5 wt% to prepare the PLC solution. After evaporation of the 1-decanol in an oven at 80 °C for 10 min or in room temperature for 2 h, we embedded the LMPs circuit with the PLC solution. After evaporation of $CH_2Cl_2$ at room temperature for 12 h, we peeled the MPC-PLC membrane off the PET substrate.

#### Characterization of the MPC-PLC membrane

We observed the morphologies of the liquid metal particles and the MPC-PLC membrane by using scanning electron microscopy (SEM, SU8220, Hitachi, Japan) and transmission electron microscope (TEM, T20, FEI, US). The LMPs inside the PLC membrane were fully degraded and removed by adding excessive hydrochloric acid (Beijing Chemicals Works, China) and rinsing by ultrapure water (Thermo Fisher Scientific, US) over 3 times. We analyzed the diameter of the liquid metal particles by using the ImageJ (NIH, US). We bent the MPC-PLC membrane for 180° for 1000 cycles and measured the conductivity after each 250 cycles and normalized the resistance value to the initial value. For the degradation test, we incubated the MPC-PLC membrane (n= 48) in the PBS solution (pH=7.4, 37°C, 5% $CO_2$) for 8 weeks and refreshed the PBS solution every day. In the first week, we took out 3 samples every day and afterwards we took out 3 samples every week, washed them with ultrapure water for over 3 times, removed the residual water by using a lyophilizer (FD-1A-50, Biocool, China) and tested the weight and analyzed the mass loss.

**Permeability of MPC-PLC membrane**

We used a 70 μm MPC-PLC membrane, folded in half to wrap the 2 ml solution which contained 2.4 mg/ml $Mg^{2+}$, 2 mg/ml $Fe^{3+}$, and 2.3 mg/ml $Ca^{2+}$, as the sample to measure the permeability of these cations. The whole part was sealed by dialysis clamps and put in a beaker filled with 400 ml deionized water for 18 days. We measured the concentrations of these cations in the beaker at 2, 5, 10, 18 day by inductively coupled plasma mass spectrometry (ICP-MS, Agilent 7700x, US), and set the initial deionized water as the control group.

**Preparation of the MPC-PLC electronic blood vessel**

We prepared the MPC-PLC electronic blood vessel by rolling up the MPC-PLC membrane with the assistance of a PTFE mandrel (Taobao, China). The diameter of the PTFE mandrel is 1.8 mm. We used a biomedical fibrin glue (Fibrin Sealant Kit, Puji Medical Technology Development Co., Ltd, Hangzhou, China) to facilitate the combination of different layers.

**Cell patterning**

We sterilized the MPC-PLC membrane by radiation with a cobalt radiation device (Co 60, 10-130 Gy $min^{-1}$, Peking University, China). Before cell seeding, the MPC-PLC membrane was incubated with fibronectin solution (50 μg $ml^{-1}$) for 6 h at room temperature to facilitate the cell attachment. We deployed a PDMS chip with three channels to seed three kind of blood vessel cells, human umbilical vein endothelial cells (HUVECs, ATCC, US), smooth muscle cells (SMCs, ATCC, US), and fibroblasts (ScienCell, US), on the surface of the MPC-PLC membrane, respectively. After overnight incubation, we removed the PDMS chip off the cell-laden MPC-PLC membrane and rolled the cell-laden MPC-PLC membrane into a natural blood vesselmimicking tubular structure with the assistance of a PTFE mandrel. Three kind of blood vessel cells were distributed in different layers of the electronic blood vessel sequentially, i.e., HUVECs (inner layer), SMCs (middle layer), fibroblasts (outer layer). To deliver different cells to different layers accurately, we designed the width of each channel according to the circumferences of corresponding layers. We cultured the cell-laden MPC-PLC membrane and the MPC-PLC electronic blood vessel in DMEM supplemented with 10% fetal bovine serum (5% $CO_2$, 37 $^oC$). To image the cells, we stained the HUVECs (blue), SMCs (green), fibroblasts (red) by CellTracker violet, green, deep red (Life Technologies, US), respectively. We used a confocal laser scanning microscopy (CLSM, LSM710, Zeiss, Germany) to capture the fluorescent images.

**Cytotoxicity of the MPC-PLC electronic blood vessel**

We cultured the MPC-PLC electronic blood vessel for 2 weeks. We used Calcein-AM green to test the viability of the cells embedded in the MPC-PLC electronic blood vessel. We washed the cell-laden electronic blood vessel with fresh PBS solution for 3 times and incubated in Calcein-AM green solution at a concentration of 5 μL $mL^{-1}$ for 20 min. After 3 times wash with PBS, we fixed the cells with 4% paraformaldehyde aqueous solution for 10 min. We used confocal laser scanning microscopy (CLSM, LSM710, Zeiss, Germany) to capture the fluorescent images. We took the multi-layered images of the sample and performed 3D reconstruction of the cells by using the ZEN software (Zeiss, Germany). We re-spread the electronic blood vessel before taking image.

**Hemolysis test**

We extracted the fresh rabbit blood into the anticoagulant tube and centrifuged at 1500 rpm for 10 min. We collected the erythrocytes after 3 times washing with saline. We prepared liquid metal particles in saline at different concentrations, 5 mg PLC membrane, and 5 mg PLC-MPC membrane. We mixed the samples with erythrocytes. The final hematocrit level of red blood cell is about 4%. After 4 h incubation at 37 $^oC$, we extracted the supernatant after centrifugation at 12000 rpm for 10 min and measured the absorbance at 540 nm by the UV-Vis analysis. We used saline as the negative control, and pure water as the positive control.

**Cell proliferation by *in vitro* electrical stimulation**

To evaluate the proliferation by electrical stimulation, we patterned HUVECs on the MPC-PLC elctronic blood vessel and incubated overnight in DMEM supplemented with 10% fetal bovine serum (5% $CO_2$, 37 $^oC$). We connected the MPC-PLC electronic blood vessel to a multi-channel electrochemical station (1040C, CHI, China) to generate the direct current. We connected six samples to separate channels with different voltage output, setting a sample without electrical stimulation as control. We generated the electrical field of 25 mV $mm^{-1}$, 50 mV $mm^{-1}$, 75 mV $mm^{-1}$, 100 mV $mm^{-1}$, 200 mV $mm^{-1}$, 400 mV $mm^{-1}$, by exerting a voltage of 25 mV, 50 mV, 75 mV, 100 mV, 200 mV, 400 mV, respectively. The interval between each electrode is 1 mm. After connecting to the electrochemical station, we incubated the cell-laden electronic blood vessel for 2 days in DMEM supplemented with 10% fetal bovine serum (5% $CO_2$, 37 $^oC$). We stained nuclei (blue) with Hoechst 33342 (Invitrogen, USA) and stained the living and dead cells with Calcein-AM (green) and PI (red), respectively. We observed the HUVEC proliferation by a confocal laser scanning microscopy (CLSM, LSM710, Zeiss, Germany) and analyze the image by ImageJ. We used the CCK-8 kit (Dojindo, Japan) to test the cell viability. We digested cells with 0.25% trypsin (Thermo fisher, US) from the substrate and cultured them in a 96-well plate for 6 h. We measured the absorbance at 450 nm by using a microplate reader. We analyzed the data by using Graphpad Prism 8.

**Cell migration by *in vitro* electrical stimulation**

We developed a wound healing model to evaluate cell migration by *in vitro* stimulation. We patterned HUVECs on a PDMS substrate and made a scratch on the cell-laden PDMS substrate by using a 10 μl tip. We attached the unfolded electronic blood vessel onto the cell-laden PDMS substrate. We connected the electronic blood vessel to a multi-channel electrochemical station (1040C, CHI, China) to generate the direct current. We used the same parameters as in the proliferation experiment. After 22 h incubation and electrical stimulation, we stained the HUVECs with DiO dye (Life Technologies, US) and observed the HUVEC migration by a confocal laser scanning microscopy (CLSM, LSM710, Zeiss, Germany) and analyze the image by ImageJ.

**Lyophilization of GFP plasmid DNA on the MPC-PLC membrane**

We lyophilized the GFP plasmid DNA onto the sterilized MPC-PCL membrane before seeding cells. We added a PDMS chip on top of the MPC-PLC membrane and added 2 ml of the GFP plasmid DNA solution at a concentration of 40 μg ml$^{-1}$ before transferring into a lyophilizer (LGD-0.1, Shanghai Kanxin Instrument Company, China). The GFP plasmid DNA was dehydrated and fixed on the surface of the MPC-PLC membrane after overnight lyophilization.

***In vitro* GFP plasmid DNA delivery via electroporation**

We connected the cell-laden MPC-PLC electronic blood vessel with an electroporator (Electro Square Porator TM ECM 830, BTX, USA). Before electroporation, we washed the samples with PBS solution 3 times and immersed the cell-laden blood vessel into the GFP plasmid DNA solution (RiboBio, China) at a concentration of 40 μg ml$^{-1}$ for 10 min. We applied 5 electrical pulses by exerting a square wave pulse. The voltage is 60 V, the pulse duration is 100 μs, and the pulse interval is 1 s. After electroporation, we disconnected the electronic blood vessel with the electroporator and incubated it for 2 days in DMEM supplemented with 10% fetal bovine serum (5% $CO_2$, 37 °C). We re-spread the electronic blood vessel and fixed the cells with 4% paraformaldehyde aqueous solution for 10 min and performed cytoskeleton staining. After 3 times of rinses with fresh PBS, we treated it with 0.1% TritonX-100 solution for 10 min. After 3 times of rinses with fresh PBS, we treated it with 3% bovine serum albumin (BSA) to avoid the non-specific binding activity. We stained the nucleus and F-acting with Hoechst 33342 (Invitrogen, US) solution (1:1000 dilution in PBS) for 5 min and Alexa Fluor 488 Phalloidin (Invitrogen, US) solution (1:200 dilution in PBS) for 20 min, respectively. Before imaging, we rinsed the sample for 5 times with PBS solution and mounted with glycerin solution (70% wt in PBS). We used confocal laser scanning microscopy (CLSM, LSM710, Zeiss, Germany) to capture the fluorescent images.

**Electroporation on isolated rabbit vascular tissues**

We dissected a 3 cm vascular tissue from the rabbit carotid artery and cut it longitudinally to flatten the tissue. We attached the MPC-PLC membrane lyophilized with the GFP plasmid DNA onto the tissue. After the plasmid DNA dissolved in the fluids, we applied 5 electrical pulses by exerting a square wave pulse. The voltage is 60 V, the pulse duration is 100 μs, and the pulse interval is 1 s. After 2 d incubation in the DMEM supplemented with 10% fetal bovine serum (5% $CO_2$, 37 °C), we stained the nuclei by DAPI and observed the GFP expression with confocal laser scanning microscopy (CLSM, LSM710, Zeiss, Germany).

**Mechanical test of the MPC-PLC blood vessel**

**Stress-strain test**

We preformed the stress-strain test of the MPC-PLC electronic blood vessels, setting the native carotid artery as a control. We used a universal tensile test machine (Instron 3365, US) to perform the test. We recorded the stress-strain data by Instron Bluehill software. The total length of the tested MPC-PLC electronic blood vessels and native carotid arteries is 30 mm. The gauge length is 15 mm. The drawing speed is set at 10 mm/min until failure. We processed the data and calculated the ultimate tensile strength, modulus, and strain at break by using Origin Pro. 5 individual MPC-PLC electronic blood vessels and 3 individual native carotid arteries were tested.

**Burst pressure and compliance**

We performed the burst pressure and compliance tests by using a home-made perfusion system consisting of a pressure gage (AZ 82100 and AZ 8205, Taiwan) and peristaltic pump (PhdUltra, Havard Apparatus, US). We connected two ends of the

MPC-PLC electronic blood vessels or native carotid arteries with two individual PE tubes. We connected the PE tubes with the perfusion system to form a loop. We controlled the pressure inside the loop by adjusting the velocity of flow. We took images of electronic blood vessels with changing diameters according to the pressure change and recorded the burst pressure. 5 individual MPC-PLC electronic blood vessels and 3 individual native carotid arteries were tested. We used ImageJ (NIH, US) and Prism (Graphpad, US) to process the images and measure the diameter changes. We calculated the compliance (C) by following formula:

$$D_{inner} = D_{outer} - h_{wall}$$

Where $D_{inner}$ and $D_{outer}$ represent the inner and outer diameter of the blood vessels respectively, $h_{wall}$ represents the thickness of the wall of the blood vessels.

$$\mathrm{C} = \frac{\frac{D_{inner}(P_2) - D_{inner}(P_1)}{D_{inner}(P_1)}}{P_2 - P_1} \times 10^4$$

Where $P_1$ and $P_2$ represent the lower and higher pressure, $D_{inner}(P_1)$ and $D_{inner}(P_2)$ represent the inner diameter in the pressure of $P_1$ and $P_2$ respectively. The compliance is expressed as percent diameter change per 100 mmHg.

***In vivo* study**

We conducted the rabbit study in the Center for Cardiovascular Experimental Study and Evaluation in Fuwai Cardiovascular Hospital (FCH, Beijing, China). All the rabbit studies were carried out in compliance with the protocols approved by Institutional Animal Care and Use Committee at the Center for Cardiovascular Experimental Study and Evaluation of the FCH. 8 rabbits were permitted by the ethics committee to be used as an early stage study. We used New Zealand rabbits (age: 200-300 days, body weight: 3-4 kg) in this study. We performed an end-to-end anastomosis procedure to implant the MPC-PLC blood vessel on carotid arteries of rabbits (n=6). We set two rabbits as a blank control group. We anesthetized the rabbits by injecting 1 ml 3% (3g 100 $ml^{-1}$) pentobarbital sodium solution from the auricular vein. The rabbits were connected to the active breathing control system (Primus,

Dräger, Germany) and life monitor system (LifeWindow 6000, Digicare, US), which provided automated anesthesia and ensured the active ventilation and *in situ* monitoring. The hair around the neck of rabbits was removed after sterilization. We incised the epidermal layer to expose the left internal carotid artery. We cross-clamped the proximal and distal ends of the carotid artery with two hemostatic clamps, transected a 1.5 cm segment, and performed end-to-end anastomosis of the electronic blood vessel to the carotid artery (2 cm in length, 2 mm in diameter) by a 8-0 suture (Polypropylene, PROLENE BLUE, Ethicon, US). After implantation, we tested the blood flow within the MPC-PLC electronic blood vessel by using perivascular flow module (TS410 &420, Transonic, US), which is a gold standard for animal blood flow measurement after implantation. After confirmed that the blood flow is strong and smooth, we closed the wound by using 7-0 suture (Polypropylene, Surgipro™, COVIDIEN, Ireland). After operation, the rabbits were treated with penicillin (800 thousand doses per rabbit per day) for 3 days and fed normally. No heparin or any other anticoagulant was used before, during, or after implantation procedure.

**Doppler ultrasound imaging**

We performed Doppler ultrasound imaging for all the rabbits every two weeks for 3 months. We used the Vivid E9 Ultrasound system (GE Healthcare, Norway) and a vascular imaging transducer to obtain the 2D and 4D images and videos of the electronic blood vessels. We obtained the color ultrasonography, synchronized pulse of the operative site, and corresponding blood flow velocity. We measured the diameter of the electronic blood vessels from different directions. All the results were saved as pictures and videos.

**Arteriography**

We performed arteriography for all the rabbits every month for 3 months. We anesthetized the rabbits by injecting 1 ml 3% (3g 100 $ml^{-1}$) pentobarbital sodium solution from the auricular vein. The rabbits were connected to the active breathing control system (Primus, Dräger, Germany) and life monitor system (LifeWindow

6000, Digicare, US), which provided automated anesthesia and ensured the active ventilation and *in situ* monitoring. We used a cardiovascular and interventional imaging system (Innova 2100-IQ, GE Medical Systems SCS, GE Healthcare, France) to diagnose the patency of the MPC-PLC electronic blood vessels. Iopromide contrast media (Ultravist 370) was injected into the vascular system to visualize the blood vessel and blood flow in real time via digital subtraction angiography. The electronic blood vessel exhibited auto-radiographic under the arteriography due to the existence of LMPs based circuits. All the diagnosis results were saved as pictures and videos.

**Blood test**

Followed by the ultrasound imaging at each time points, we extracted 3 ml blood sample from the rabbits for inflammation and infection tests, including blood routine examination, biochemical tests and ELISA assay of three important proteins, i.e., interleukin-6 (IL-6), procalcitonin (PCT), and C-reactive protein (CRP). 2 ml blood was sent to the Fuwai hospital for the blood routine examination and biochemical tests. 1 ml blood was processed in the lab for ELISA tests. We collected all the samples and tested them in the end time point by using three rabbit ELISA kits (Rabbit IL-6 ELISA kit, PCT ELISA kit, and CRP ELISA kit, MyBioSource, US).

***Ex vivo* study**

We explanted all the MPC-PLC electronic blood vessels and their major organs, including heart, liver, spleen, lung, and kidney, 3-month post-implantation. In the meantime, we also sacrificed two normal rabbit in the blank control group to obtain the native carotid artery and their major organs. We preserved all the samples in 4% paraformaldehyde aqueous solution separately. Part of the samples were dehydrated by using an automatic tissue dehydrating machine (ASP200s, Leica, Germany) and paraffin-embed by using a paraffin-embedding machine (EG1150 System, Leica, Germany). We cut the samples into 6 μm thick sections by using a microtome (RM2235, Leica, Germany) and de-paraffinized the samples twice (10 min per time) with dimethylbenzene and serially rinsed 100% ethanol, 90% ethanol, 80% ethanol,

70% ethanol, and deionized water for 5 min each step. We stained the cross sections of the MPC-PLC electronic blood vessels and the native carotid arteries with hematoxylin and eosin (H&E), Masson's trichrome, and Verhoeff's and affixed the stained sections to coverslips with Gelvatol mounting media. We stained the cross sections of the major organs from MPC-PLC electronic blood vessels implanted group and the control group with hematoxylin and eosin (H&E), Masson's trichrome and affixed the stained sections to coverslips with Gelvatol mounting media. We captured the images by using an upright microscope (DM4000M, Leica, Germany). All the images are representative of at least 3 independent samples.

**Statistical analysis**

We conducted image analysis of liquid metal particles, MPC-PLC membrane, mechanical tests data, and doppler ultrasound data by using ImageJ (NIH, US), Prism 8 (Graphpad, US). We recorded the stress-strain data by using Instron Bluehill software (Instron, US). All the statistical data are expressed as mean ± standard deviation with a group number n described in the caption.

### Supplemental Figures

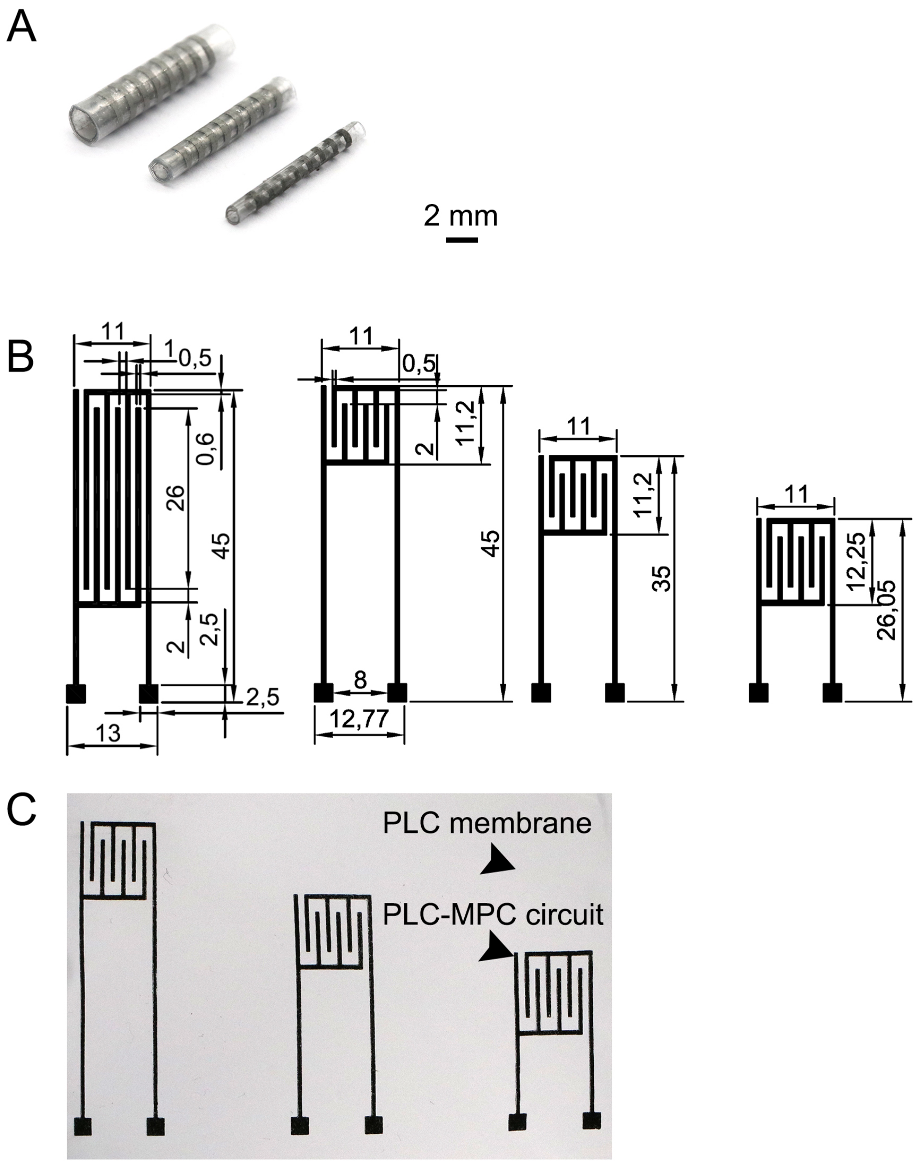


**Figure S1. Electronic blood vessels with different circuit designs and with different diameters. (A)** Electronic blood vessels with diameter of 2mm, 1 mm, 0.5 mm, respectively. **(B)** MPC-PLC membranes with different circuit designs targeting different blood vessel layers. Numbers are in mm. **(C)** Snapshots of the MPC-PLC membranes with different circuit designs.

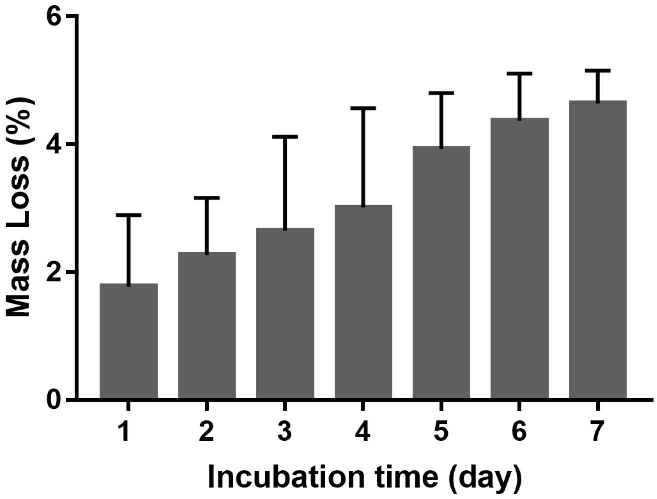


**Figure S2. Mass loss of the MPC-PLC membrane during the first week.**

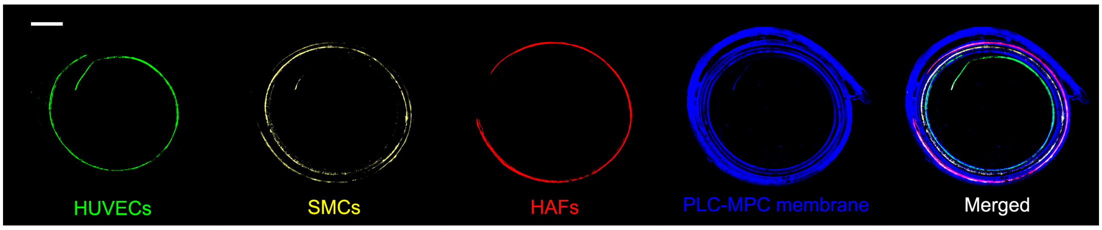


**Figure S3. Distribution of cells and the MPC-PLC membrane in the electronic blood vessel.** Green represents HUVECs, stained by CellTracker DiO; yellow represents SMCs, stained by CellTracker DiI; red represents HAFs, stained by CellTracker DiD; blue represents PLC-MPC membrene, stained by CellTracker Blue. Scale bar, 500 μm.

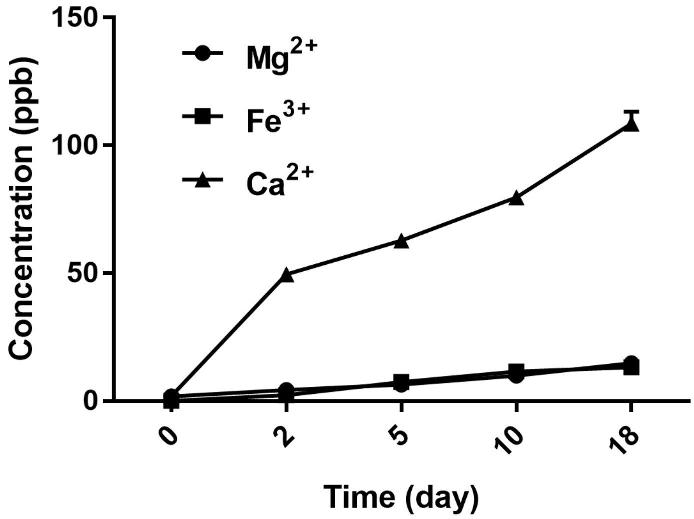


**Figure S4. Ionic permiability of the MPC-PLC membrane.**

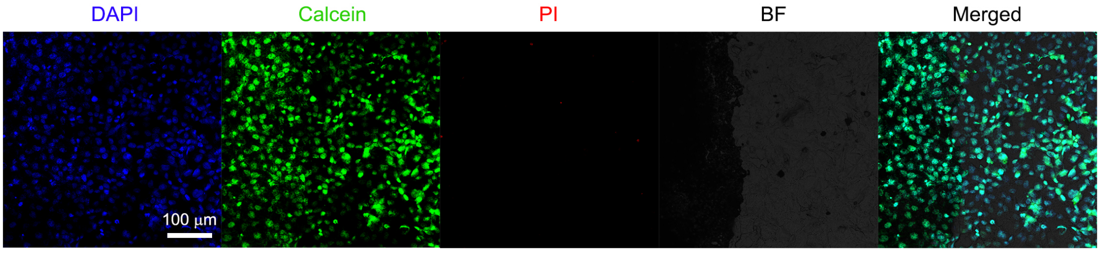


**Figure S5. Cytotoxicity test after 10-day electrical stimulation.** Blue represents nuclei, stained by DAPI; live cells were stained by CalceinAM green; dead cells were stained by PI. Scale bar, 100 µm.

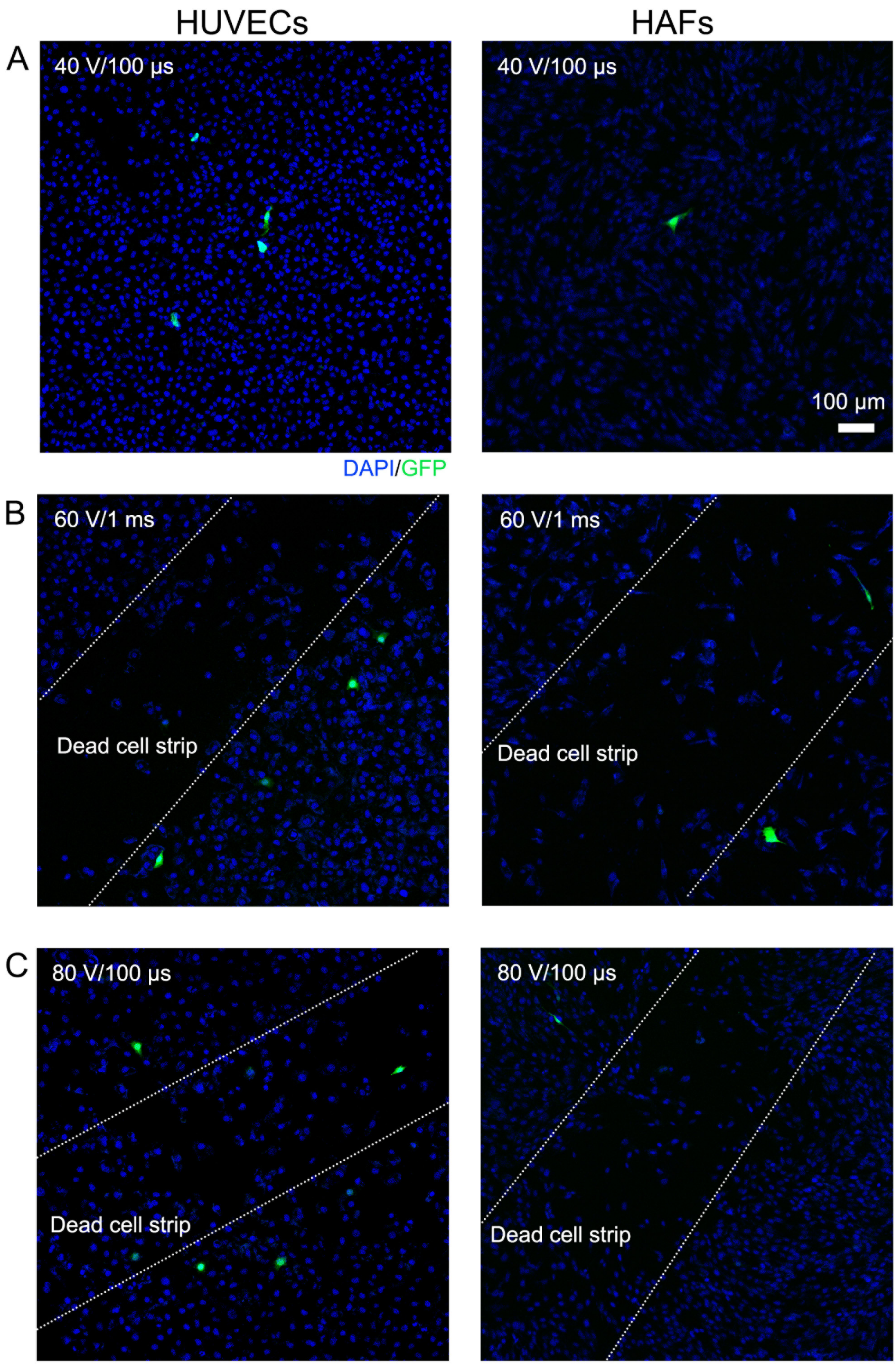


**Figure S6. Electroporation with different voltages and pulse durations.** (**A**) Electroporation under 40 V, 100 μs duration caused low efficacy or no transfection. (**B**) Electroporation under 60 V, 1 ms duration caused low efficacy and a dead cell strip. (**C**) Electroporation under 80 V, 100 μs duration caused low efficacy and a dead cell strip. Blue represents nuclei, stained by DAPI; Green represents green fluorescent protein. Scale bar, 100 μm.

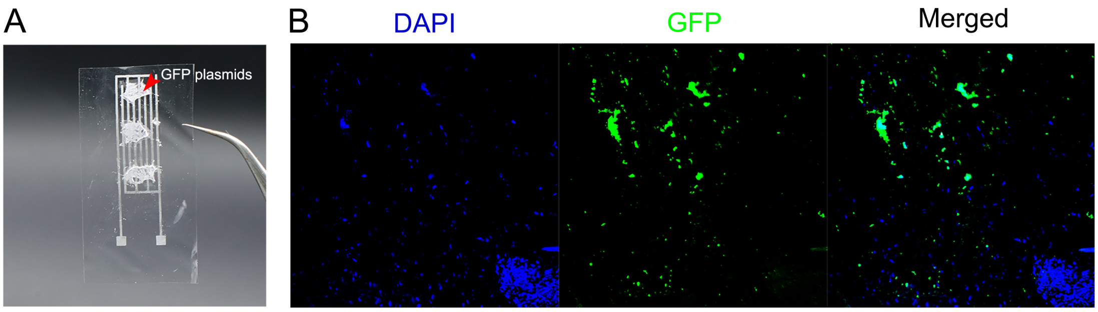


**Figure S7. Electroporation of the isolated rabbit vascular tissue.** (A) Lyophilization of GFP plasmids DNA on the MPC-PLC membrane. (B) Electroporation of the isolated rabbit vascular tissue. Blue represents nuclei; green represents GFP.

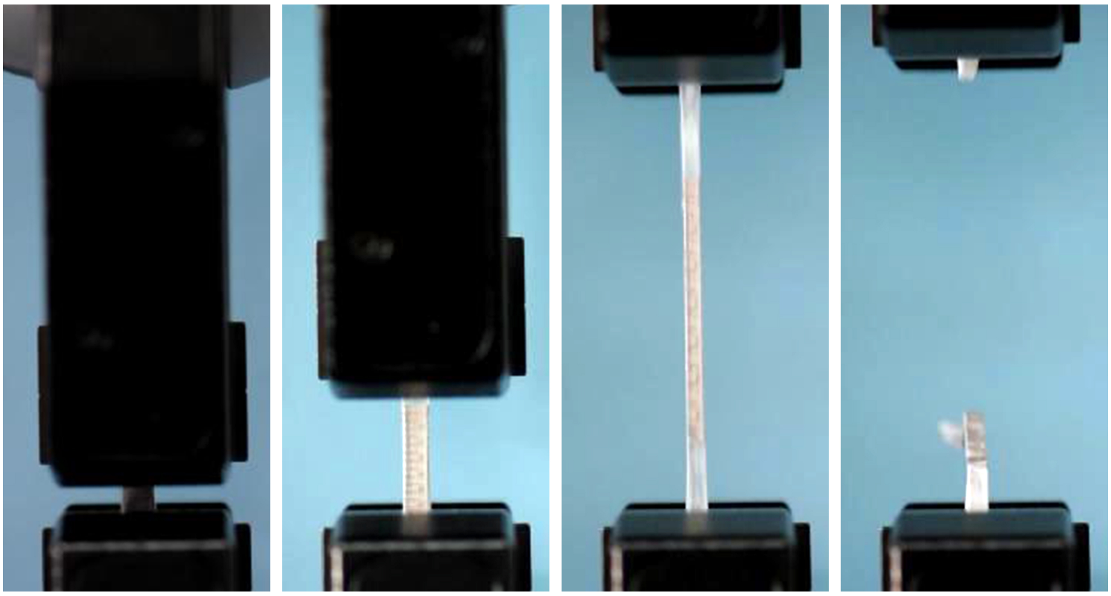

**Figure S8.** Snapshots during the tensile test of the MPC-PLC blood vessel.

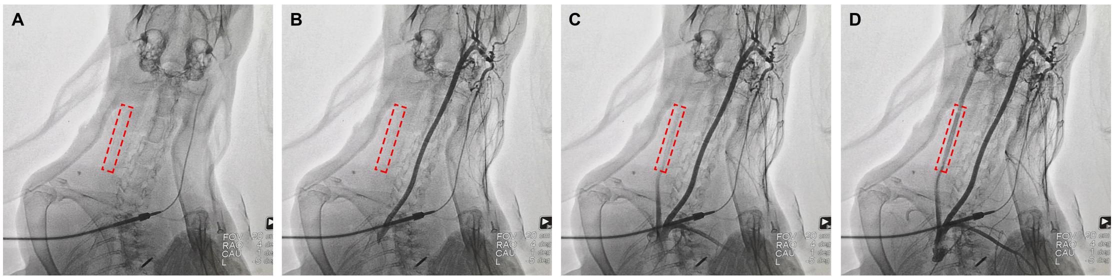


**Figure S9.** Time-serial snapshots of the arteriography.

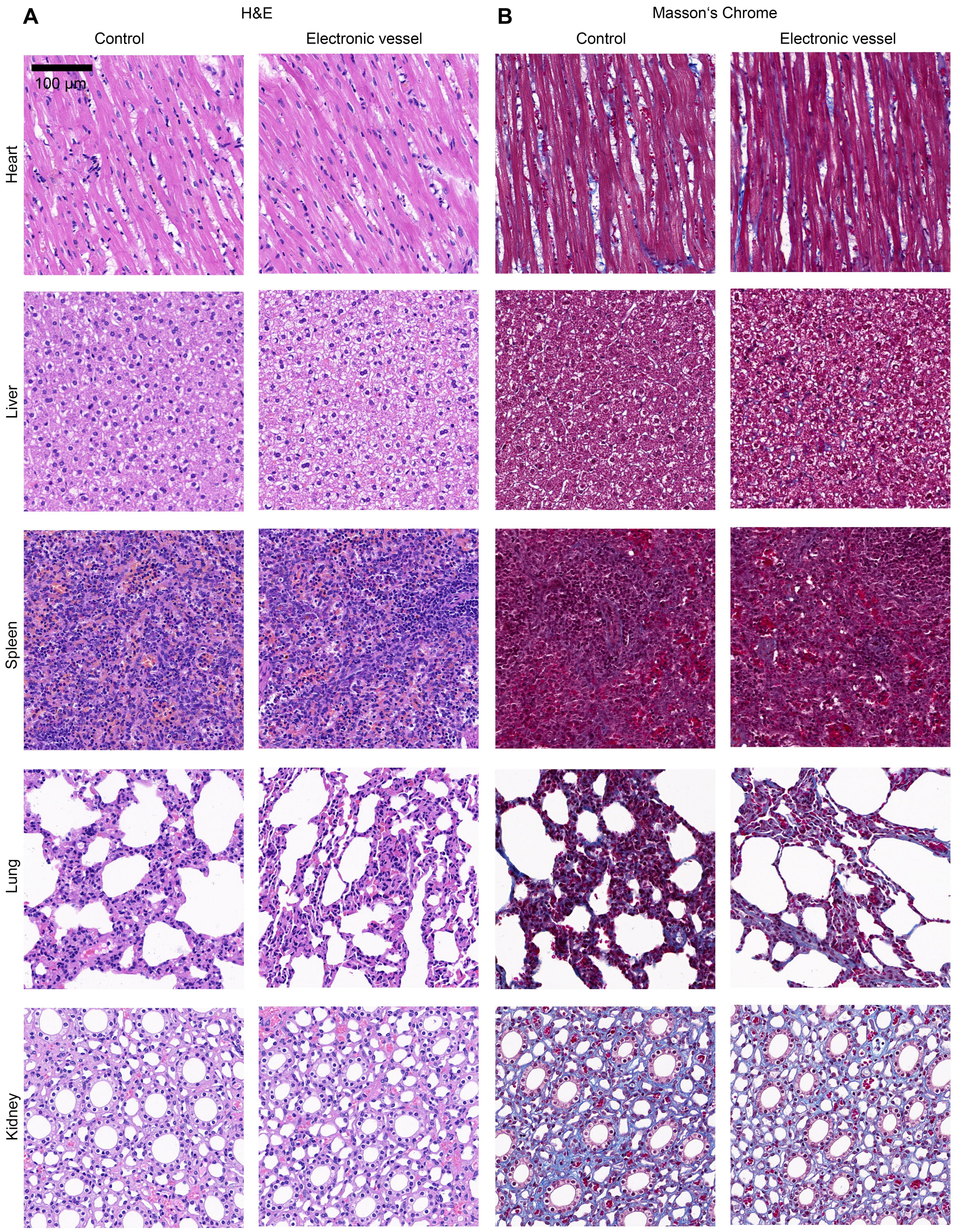


**Figure S10. *In vivo* toxicity of the electronic blood vessel. (A)** H&E staining and **(B)** Masson's Chrome staining of cross section of the major organs of the rabbits, including heart, liver, spleen, lung, kidney. Left column represents the control group, right column represents the electronic blood vessel group. There is no significant difference between two groups. Scale bar, 100 µm.

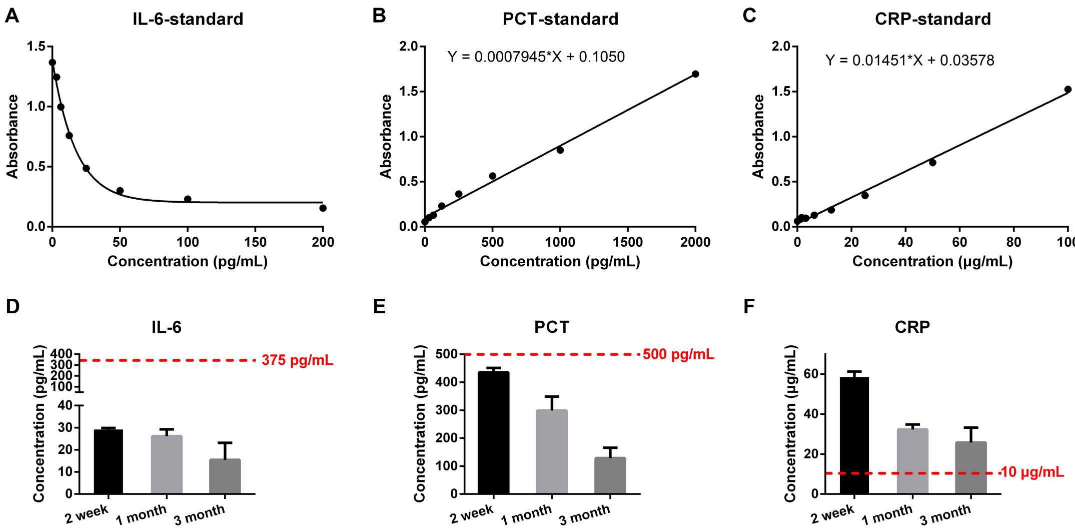


**Figure S11. Chronic inflammatory reactions after implantation. (A-C)** The standard curves of absorbance to concentration of the IL-6, PCT, and CRP. **(D-F)** The changes of the concentration of IL-6, PCT, and CRP in the serum of rabbits after implantation of the electronic blood vessel. The red dotted lines indicated the normal value of healthy rabbits.

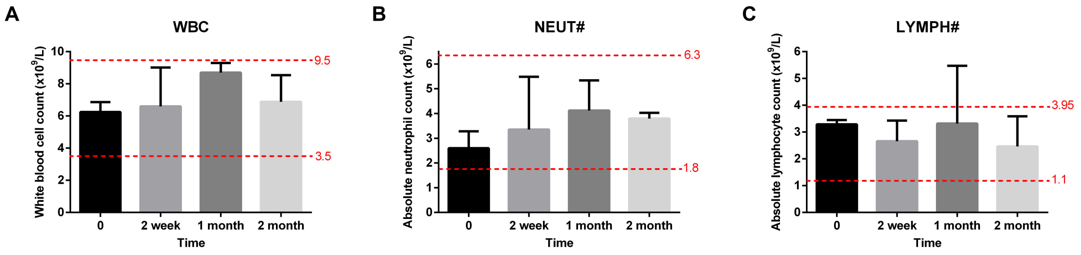


**Figure S12. Complete blood count. (A)** White blood cell count. **(B)** Absolute neutrophil count. **(C)** Absolute lymphocyte count.

### Supplemental Videos

**Video S1. Doppler ultrasound imaging 3 months post-implantation.** The electronic blood vessel allowed for good blood flow.

**Video S2. Arteriography 3 months post-implantation.** The electronic blood vessel matched with the native carotid artery very well and allowed for excellent blood flow.
